# A Method to sort heterogenous cell populations based on migration in 2D and 3D environments

**DOI:** 10.1101/2020.05.06.080234

**Authors:** Aditya Arora, Jorge Luis Galeano Niño, Myint Zu Myaing, Bakya Arasi, Ruby Yun-Ju Huang, Ramanuj Dasgupta, Maté Biro, Virgile Viasnoff

## Abstract

Whereas phenotypic assays such as Boyden chambers and wound healing assays can easily be employed to characterize the migratory potential of cells at the population level, few methods exist that can sort subpopulations of cells based on their migratory behaviour from an initial heterogeneous pool. In this paper, we present an approach to sort migratory cancer and immune cells based on their spontaneous migration in 2D and 3D microenvironments. Using this method, which is easy to implement and readily scalable, millions of live cells can be sorted based on their migratory characteristics and then subjected to downstream genomic, molecular and functional tests. We reveal that enrichment of the most migratory cytotoxic T lymphocytes yields a pool of cells with enhanced cytotoxicity against cancer cells. This new functional sorting method opens new avenues for the precise characterization of the mechanisms underlying hitherto unexplained heterogeneities in migratory phenotypes within a cell population, and for the targeted enrichment of the most potent migratory leukocytes in immunotherapies.

## INTRODUCTION

Cell migration plays a pivotal role in all stages of the life of a multicellular organism. During development, cells migrate over long distances to give rise to tissue morphogenesis. Similarly, cell migration is crucial in wound healing both for supply of progenitor cells and immune cells to the site of injury which enable regeneration and prevent infections respectively. Further, aberrant migration of diseased cells such as cancer cells leads to their dissemination and thus, facilitates metastasis. As a result, a variety of methods have been developed to study the migration of cells. Migration driven by chemotaxis and transmigration are often tested using Boyden chambers ^1^, wherein migratory cells are allowed to migrate from one compartment to another through a porous membrane. Chemotactic gradients are created either by addition of soluble factors or by culture of secretory cells in one of the compartments. The detection of the migrated cells is usually spectrophotometric using a plate reader or via flow cytometry. Wound healing assays (also referred to as scratch assays) are preferentially used to study collective cell migration. They consist in creating a gap or a scratch in a two-dimensional (2D) monolayer of cells and to image the collective invasion of the freed space by the surrounding cells. Time lapse imaging allows basic measurements such as the speed of gap closure often considered as a quantitative proxy for cell migration ability ^2^. Other quantitative measurements can be performed such as the roughness of the migration front, indicative of the cohesion in the collective behavior of the cells. This type of assay is widely used as a simple first phenotypic characterization of cancer cells. Its simplicity and straightforward adaptability to quantitative high-resolution imaging makes it an assay of choice to understand the molecular basis of collective cell migration. Approaches to study cell migration in 3D are less popular due to the technical challenges of imaging. They often rely on the generation of spheroids in a matrix environment and leaving the cell “sprout” off the initial cell cluster. Quantitative measurement of the number of sprouting cells and mean distance traversed reveals the migration potential of the tested cell population ^3^.

Frequently, these assays reveal heterogeneity of migratory behavior within a cell population, however, the quantitative analysis usually provides only a migration index averaged over the entire cell population ^4^. Despite the heterogeneity observed by imaging, these assays do not provide easy (if any) means to sort and to retrieve cells within the population based on their migration behaviours. Single cell transcriptomics now allows for quantitative assessment of the single cell heterogeneity in cancer cell populations and can reveal the drift of subpopulations of cells with enhanced metastatic potential. It thus creates a need for phenotypic migration assays that can (i) sort cells based on migration potential, (ii) retrieve easily the sorted cells in large quantity (of the order of 1 million cells), and (iii) permit subsequent genomic/molecular tests on the sorted live population. The isolation of such migratory cells facilitates identification of invasion signatures which can then be used for prognosis or as therapeutic targets ^5^. At the same time such an assay may be used to isolate cells with better motility for therapy such as in regenerative medicine and immunotherapy.

Very few such assays have been developed to date. When they exist, they are either of very low throughput, or difficult to implement or parallelise. Carlos et al., utilized an array of teardrop shaped adhesive protein patterns to sort cells based on motility, analogous to a thin layer chromatography setup ^6^. In another work, the ability of migrating cells to phagocytose fluorescent particles from a substrate was used to study and sort cells using fluorescence-activated cell sorting (FACS) ^7^. Taken together, these and some other methods ^8–10^ provide a tool for 2D motility-based sorting using complex micro-patterns, microfluidic devices, or FACS-based methods. However, due to the involvement of these high-end technologies the use of such methods is limited by their cost, complexity and scalability.

Further, there is a dearth of methods that can be used to sort cells based on their ability to migrate in 3D microenvironments. While 2D motility studies have provided significant insight into basic biological processes. Their use is still limited by the fact that most physiological cell migration occurs in 3D microenvironments.^3^ Wyckoff et al, used matrigel and growth factor loaded microneedles to capture migratory cells from tumors in living rats^11^, this has later been used perform gene expression and molecular analysis of a small number of cells to identify novel invasive signatures^12–14^. However, this method is limited by its use in larger *in vivo* tumors only. Moreover, certain cell types, such as leukocytes, expand in suspension culture and exhibit their characteristic migration behaviour only in 3D extracellular matrix microenvironments. Thus, for such cell types and for making the process more physiological, there is also a need to develop methods for *in vitro* 3D motility-based cell sorting.

Here we propose readily implementable methods to separate a faster migrating sub-population of cells from a heterogeneous population based on migration in 2D or 3D environments. The 2D migratory sorting relies on a layered PDMS micro-well device, which was demonstrated to be useful for separating a heterogeneous cancer cell population into a more homogenous population based on migration. The 3D sorting utilizes hierarchical hydrogel systems consisting of collagen micro-gels suspended in degradable bulk hydrogel. This method can not only separate cancer cells in 3D but can also be used for more challenging cell types such as primary leukocytes (here cytotoxic T lymphocytes (CTL)). We also characterise the sorted populations in terms of their EMT status, receptor clustering, transcriptomic data or cytotoxic function depending on the cell type.

## RESULTS

Both our 2D and 3D sorting approaches rely on similar principles that only differ by their implementation. Initially unsorted groups of cells are locally confined in a series of scattered predefined regions. The cells are then left to migrate spontaneously away from the original confinement zone onto (into) another substrate. In the 2D case the surrounding substrate is mechanically detached from the initial confinement zones, whereas, in the 3D case the separation is performed by selective matrix digestion. Cells are thus sorted by the physical separation of the pre- and post-migration substrates.

### Fabrication of 2D sorting device

Extensive description of the device is detailed in Materials and Methods and exemplified in Figure 1 and Supplementary Movie 1. Briefly, the 2D Migration Sorting Assay (2D-MSA) is first formed by laminating 3 layers of PDMS sheets: a protective layer at the top, then a collection layer, and finally a spacing layer. The thickness of the protective layer is not relevant, we found that 250 µm thick collection layers offer sufficient mechanical stability to allow easy subsequent peeling. We varied the thickness of the spacing layers to allow satisfactory sorting. Arrays of holes (300-500 um in diameter) are then perforated across the three-layer substrate using a laser cutter. The system is then assembled on a base PDMS layer. We then seeded cells on the substrate at 70-80% confluency. Cells that fall into the cavities and adhere after about 30 minutes, and subsequently the top layer is removed using lab forceps after rinsing the substrate. Cells are then allowed to migrate up the cavity walls until they reach the top collection layer initially devoid of any cells. The diameter and height of the cavity is optimized to offer maximum chance of migratory cells reaching the wall within the migration period (1 to 3 days). The spacing between the cavities have been optimised to preclude any cells migrating from one cavity to the other while keeping the highest density of wells to maximise the number of sorted cells. It ensures that cells can still migrate away from the confinement zone even if they are initially located in the center of the pit. Cell density can be varied based on the cell types. After 1 to 3 days, the layers are individually peeled off with tweezers. The collection layer is enriched with fast migrating cells while the base layer is enriched in slow migrating cells. Cells can be trypsinized immediately from the layer, leading to a collection of around 300 to 500 cells per well on the top layer. Alternatively, cells can be left to expand on the separated layer yielding around 2,000 – 4,000 cells per well. We practically implement the assay in a 6 well plate which usually contains 64 such cavities per well, thus making the total migratory cell yield from the plate of the order of 10^6^ cells.

**Figure 1:**
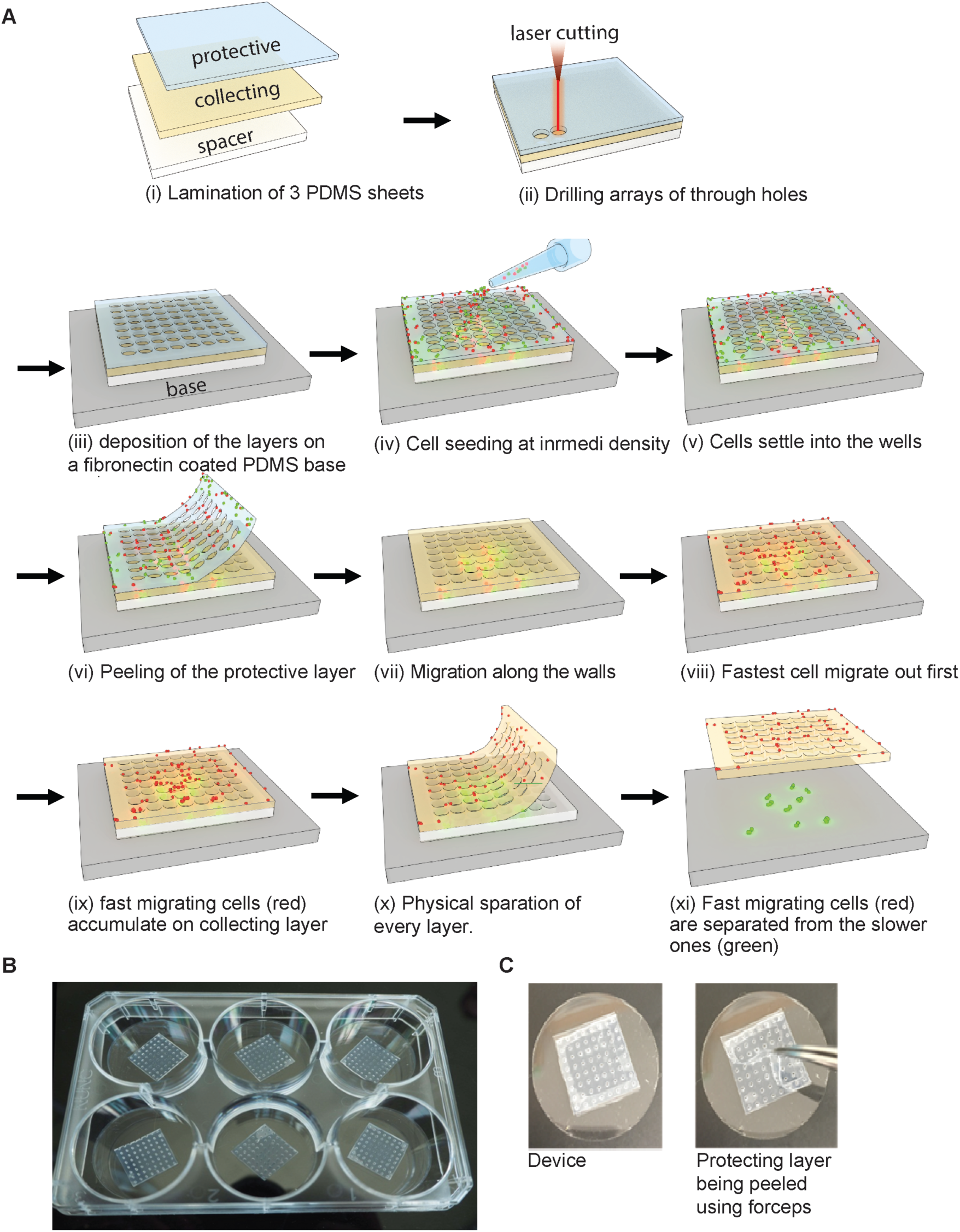
Layered-PDMS microwells for 2D migratory sorting of cells. A) Schematic depicting the process of fabrication of PDMS micro-well based device and its use for migratory sorting of cells. B) Macroscopic image of the micro-well based devices in a 6-well format. (C) Close-up of the final ready-to-use device, and the protecting layer being peeled off using forceps.

### Migratory sorting of mixture of cancer cell lines

To first validate the ability of this method to separate cells with differential motility, we used a 1:1 mixture of two established cell lines (MCF 7 and MDA MB 231) with different motility characteristics. MCF 7 and MDA MB 231 are both breast cancer derived cell lines; while the former maintains an epithelial state and lacks the ability to metastasize, the latter is largely mesenchymal with extensive migratory propensity.^15^ The two cell lines were distinguished by differentially labelling with cell tracker dyes before the experiment. The cells were seeded onto fibronectin-coated PDMS micro-well devices with spacer thicknesses of 250 and 500 µm. By the third day of incubation a fraction of cells was observed on the collection layer of the device. As expected, significantly more cells migrated out onto the collection layer in the device with a 250 µm spacer thickness (N=12). MDA MB 231 cells visibly migrated in greater abundance onto the collection layer by day 3 compared to the MCF 7 cells (Figure 2A). Quantification of this process using image analysis revealed significant enrichment of MDA MB 231 cells on the top layer of the device (Figure 2B), and revealed that these cells occupied almost fourfold higher area in the top layer versus in the bottom region of the wells. This experiment thus demonstrates that cells with higher migratory ability preferentially accumulate in the top layer of the proposed layered micro-well device.

**Figure 2:**
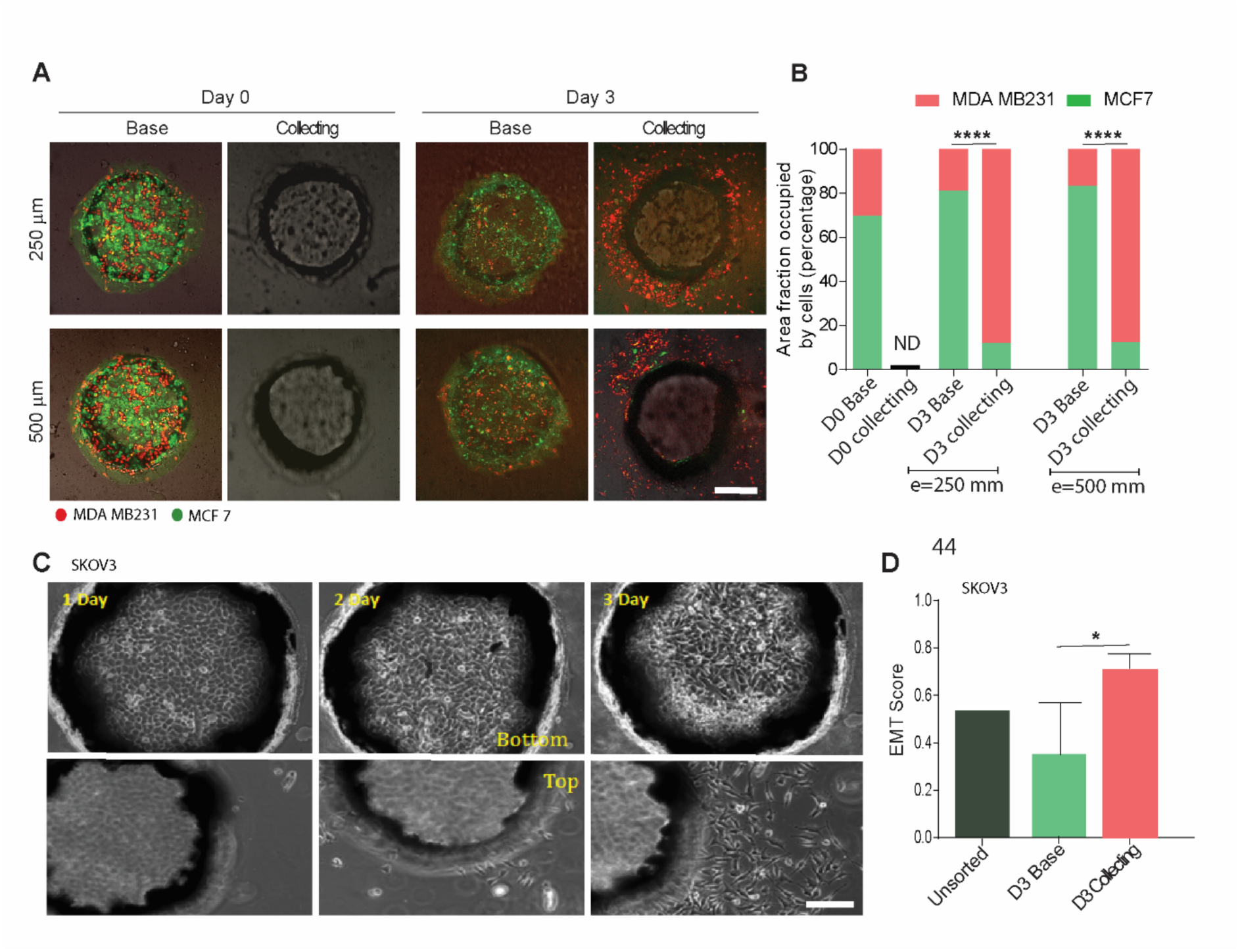
2D migratory sorting of cells with varying migratory abilities and degree of epithelial to mesenchymal transition (EMT) state. (A) Fluorescent images of bottom and top regions of microwells with MDA MB 231 (red) and MCF-7 (green) cells in microwells with different spacer height on day 1 and after 3 days of incubation (scale bar: 200 µm). (B) Quantification of area fraction occupied by MDA MB 231 and MCF 7A cells on top and bottom layers before and after migration (N = 12 microwells from four biological replicates). (C) Phase contrast images of SKOV3 cells migrating out of microwells at days 1, 2 and 3 after seeding (scale bar: 100 µm). (D) Quantification of EMT score based on expression of a panel of genes (ref) for cells isolated from top and bottom layer, secondary x axis depicts EMT score of the unsorted SKOV3 cells (N = 3). *<0.05, and ****<0.0001 by student’s T test.

### Migratory sorting of metastatic cells and correlation with Ephrin clustering and transcriptomics

We then tested the approach on head-and-neck tumour samples from two different patients. We generated Patient derived cell lines (PDCs) for each patient from lymph node metastatic sites (HN120m, HN137m). We recently demonstrated that these cells all present very distinct genomic profiles ^16,17^. They also displayed inherent heterogeneity in migration potential and Epithelial to Mesenchymal transition (EMT) scores ^18^. We used the sorting assay to segregate the fast migrating cells from the less migrating. We seeded the cells and allowed the cells to migrate either for three or five days. The cells that were allowed to migrate only for three days were subsequently allowed to proliferate for two days to maintain consistency in proliferation between the two groups. We then collected the cells and subjected them to two subsequent assays. We first used a phenotypic Ephrin A1 clustering assay to discriminate at single cell level the degree of Ephrin A2 Receptor activation using a bilayer system ^18,19^. The EphA clustering has been reported to correlate well with migration protential ^18^. EphA Scores have also been shown to be relevant to describe heterogeneity in cells at the single cell level. Figure 3 A and B shows that in the case of both the cell types (HN120M, HN137M) the cells collected on the base layer have ephrin scores that are significantly lower than scores obtained from the escaping cells after 3 days. Additionally the distribution for the unsorted cells lies between the base and collecting layer cells. Interestingly, if cells are allowed to migrate for longer (5 days), the clear distinction between EphA scores of the two cell populations vanishes (Figure 3C and D). This indicated that the first wave of cells had a distinct phenotype that could be captured if separated at an early time point and could be demonstrated using the EphA clustering.

**Figure 3:**
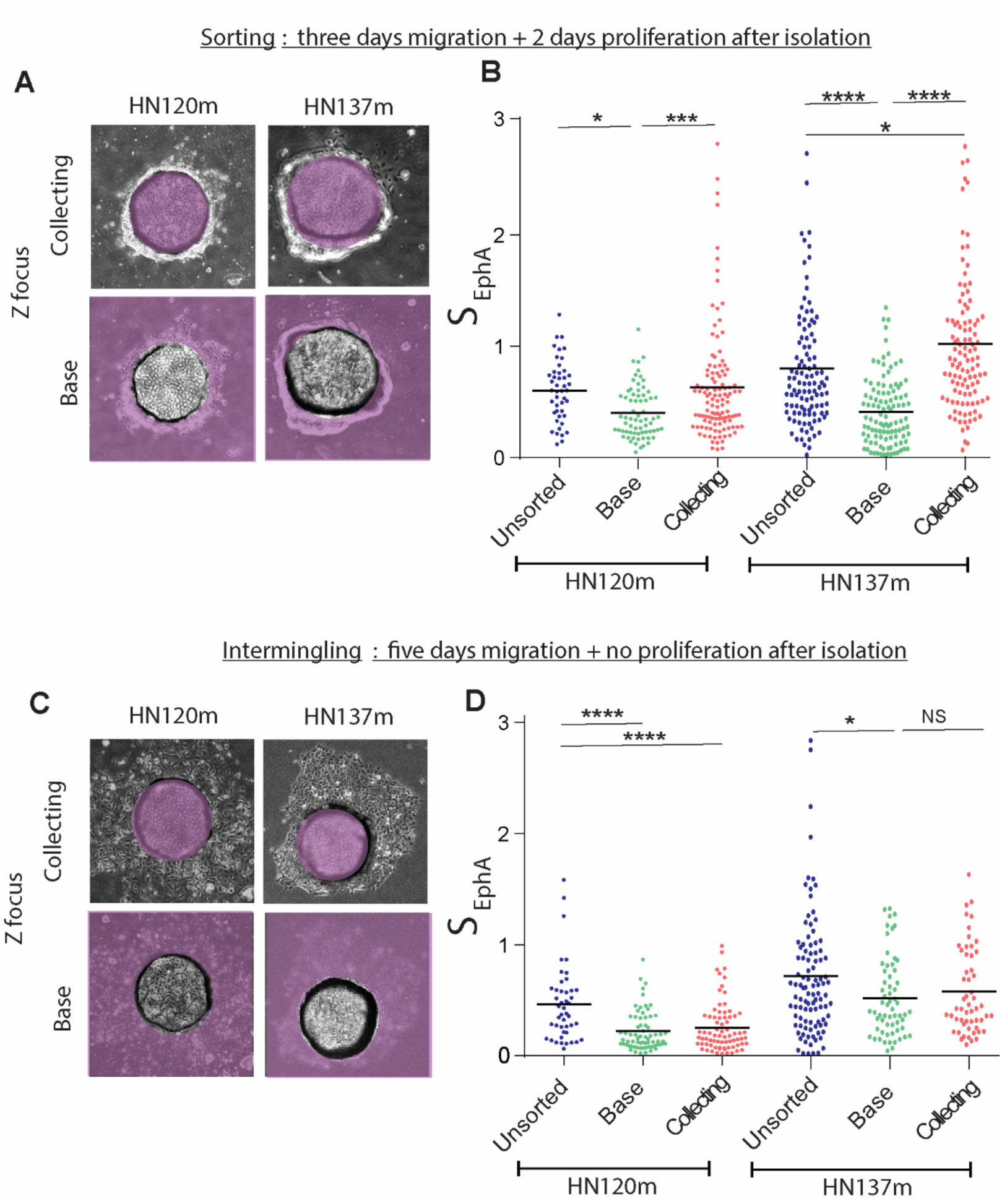
Assessment of EphA clustering heterogeneity in patient derived head and neck cancer cells using 2D migratory sorting. Phase contrast images of HN120m and HN137m cells migrating out of microwells after 3 days (A) and 5 days. Normalized Ephrin A clustering score of unsorted control, cells isolated from top and bottom layers for HN120m and HN137m cells allowed to migrate for 3 days (B) or 5 days (D). *<0.05, ***<.001, and ****<0.0001 by one-way ANOVA followed by Tukey’s multiple comparisons test. (Data points indicate the normalized score of each cell and black bars indicate the mean values, N ≥ 50 data points from three biological replicates). Purple masks cover the out-of-fucus region that does not correspond to the layer height

### Migratory sorting of cancer cells in 3D hierarchical hydrogels

We then extended the concept of entangled dissociable substrates to 3D migration. The preparation of the composite substrate is described in the Materials and Methods and Figure 4. Briefly, we first created collagen microbeads embedding cells using an emulsion in Mineral oil at 37°C. The majority of these microbeads were in the range of 100 to 200 µm in diameter, with average size being 155 µm (Figure 4B and C). The collagen microbeads are collected by gentle centrifugation, washed and resuspended in growth medium. The microbeads are then dispersed in an enzymatically degradable hyaluronic acid (HA) hydrogel mixed with very low concentration of collagen to aid cell migration ^8^. The surrounding matrix is rapidly cross-linked (<1 minute) while mixing to minimize the sedimentation of the collagen beads. The cells are left to migrate from the collagen microbeads into the surrounding cleavable matrix for 7 days. The cleavable matrix is then selectively digested using hyaluronidase enzyme for 3 hours until it liquifies to release all the migrated cells. The suspension is filtered using a cell strainer (40 µm) to separate the collagen beads containing the less mobile cells from the surrounding medium containing the most migratory cells. Each fraction can then be cultured and expanded separately. It is straightforward to scale the methods increasing the volume of the container. We typically used a set of five, 100 µl gels, and about 2-4 × 10^5^ migratory cells could be retrieved from each set.

**Figure 4:**
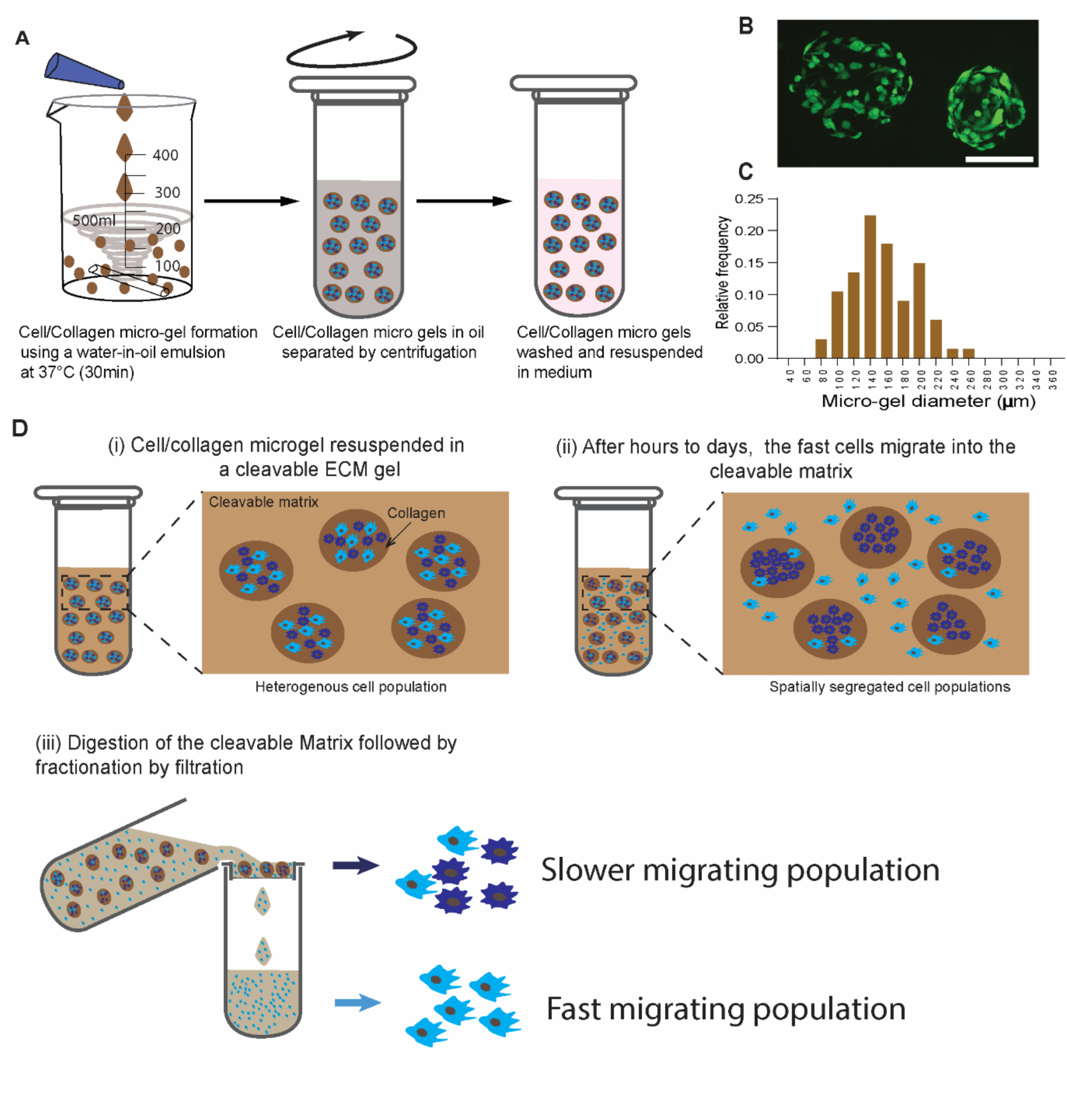
Sorting of cells based on migration through a 3D hierarchical hydrogel system. (A) Schematic depicting process for fabrication of cell laden collagen micro-gels using water-in-oil emulsion. (B) Fluorescent micrographs of MDA MB 231 GFP cells embedded in collagen micro-gels (scale bar: 100 µm). (C) Size distribution bar graph for cell laden collagen micro-gels (N=70). (D) Schematic depicting migratory sorting of cells based on differential migration through collagen micro-gels into outer hydrogel matrix.

We first validated the migration of MDA MB 231 cells from the collagen beads into the outer HA hydrogel. These experiments showed that after 7 days of incubation a large population of elongated MDA MB 231 cells were able to move across the interface of two matrices and had migrated out into the outer HA hydrogel (Figure 5A). Thereafter, we used a 1:1 mixture of epithelial MCF7 and mesenchymal MDA MB 231 cells. Figure 5B and D shows that after 7 days the population of cells in the microbead remains similar to the initial unsorted population, however the cells extracted from the surrounding HA matrix was predominantly (>95%) MDA MB231 type (Figure 5C). Taken together, we could sort and extract the fast migrating cells from the bulk population using this method.

**Figure 5:**
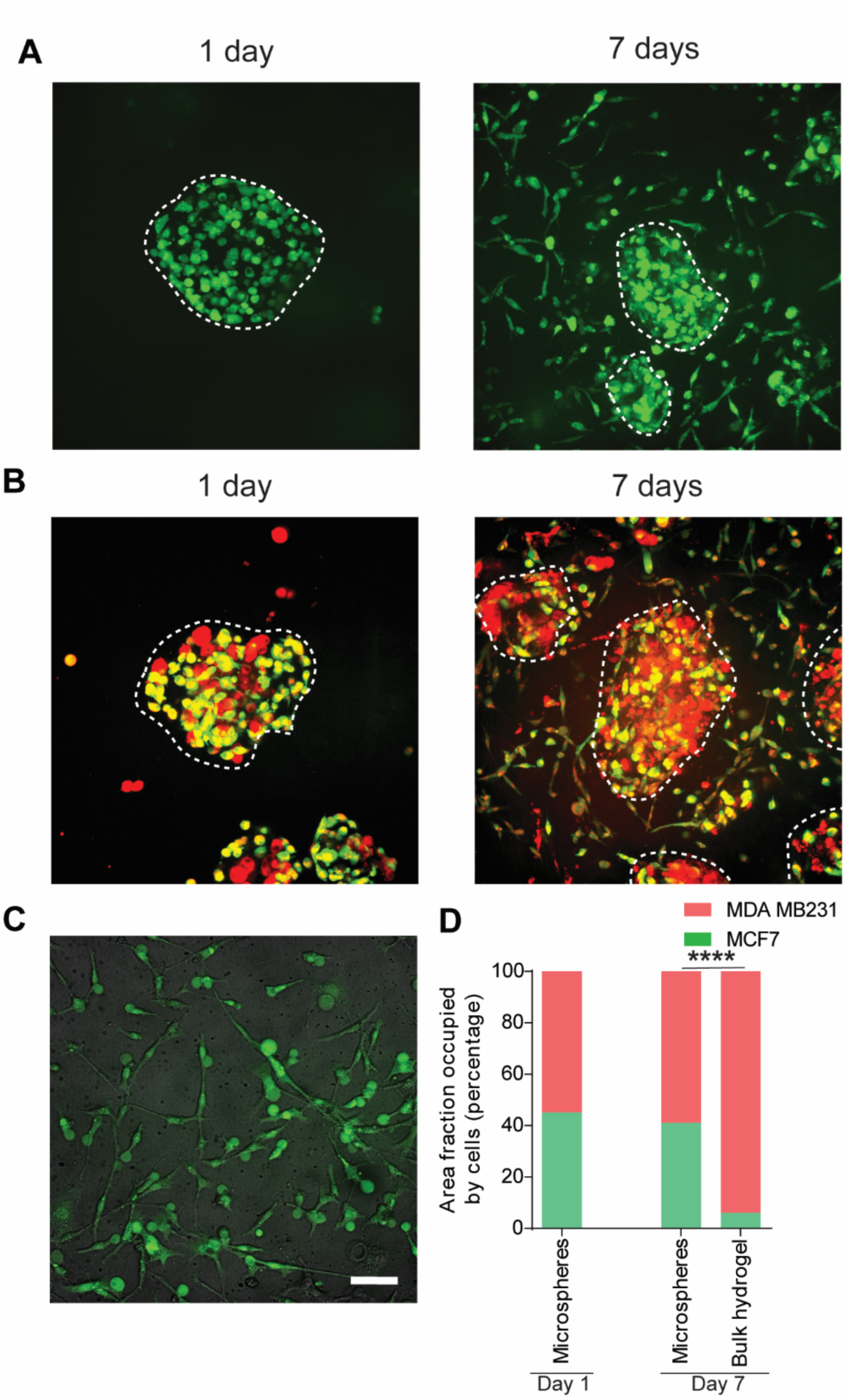
Differential migration of cells through matrices. (A) Fluorescent images of MDA MB 231 - GFP cells laden micro-gels further embedded in hyaluronic acid-collagen hydrogels post 1, and 7 days of incubation. (B) 3D migratory sorting of MDA MB 231 (labelled with GFP and cell tracker red) and MCF 7 (red) cells using the hierarchical 3D matrix enabling selective enrichment of invasive MDA MB 231 cells in the bulk hyaluronic acid-collagen hydrogel and accumulation of mixed population in the micro-gels. (C) Cells isolated after digestion of bulk hydrogel are highly enriched (~ 95%) in MDA MB 231 cells (green) (scale bar: 50 µm). (D) Quantification of area fraction occupied by MDA MB 231 and MCF 7A cells in microspheres and bulk hydrogel matrix before and after migration (N = 31 images from four biological replicates). (****<0.0001 by student’s T test)

### Migratory sorting of cytotoxic T cells and correlation with function

We then tested our approach on cells that do not readily migrate in 2D. We used primary mouse Lifeact-EGFP cytotoxic T cells ^20^. We proceeded as above for the initial embedding of the T cells in collagen beads. However, since T cells migrate much more rapidly than cancer cells and can reach speeds up to 12µm/min in collagen matrices,^21^ the experimental timeline can be markedly reduced to hours and we could therefore simply resuspend the collagen beads in cell culture medium and forgo the surrounding HA hydrogel otherwise sustains migrating cells for days. A large number of cells could emigrate from the collagen beads into the outer medium within 2 hours (Figure 6A). We then collected the cells suspended in medium and in the collagen after the 2 hour incubation period. We then measured the average migratory speed of the individual T cells from the sorted and unsorted population fractions. It reveals a clear enrichment of the fast migrating T cells in the surrounding medium, as well as a higher proportion of less migratory cells remaining in the microbeads (Figure 6B). Moreover, the migratory cell fraction had a significantly higher confinement ratio as compared to the less migratory fraction, indicating that they migrated more directionally than the retained, less migratory population (Figure 6C). Our methods thus achieved a substantial enrichment of fast migrating CTLs.

**Figure 6:**
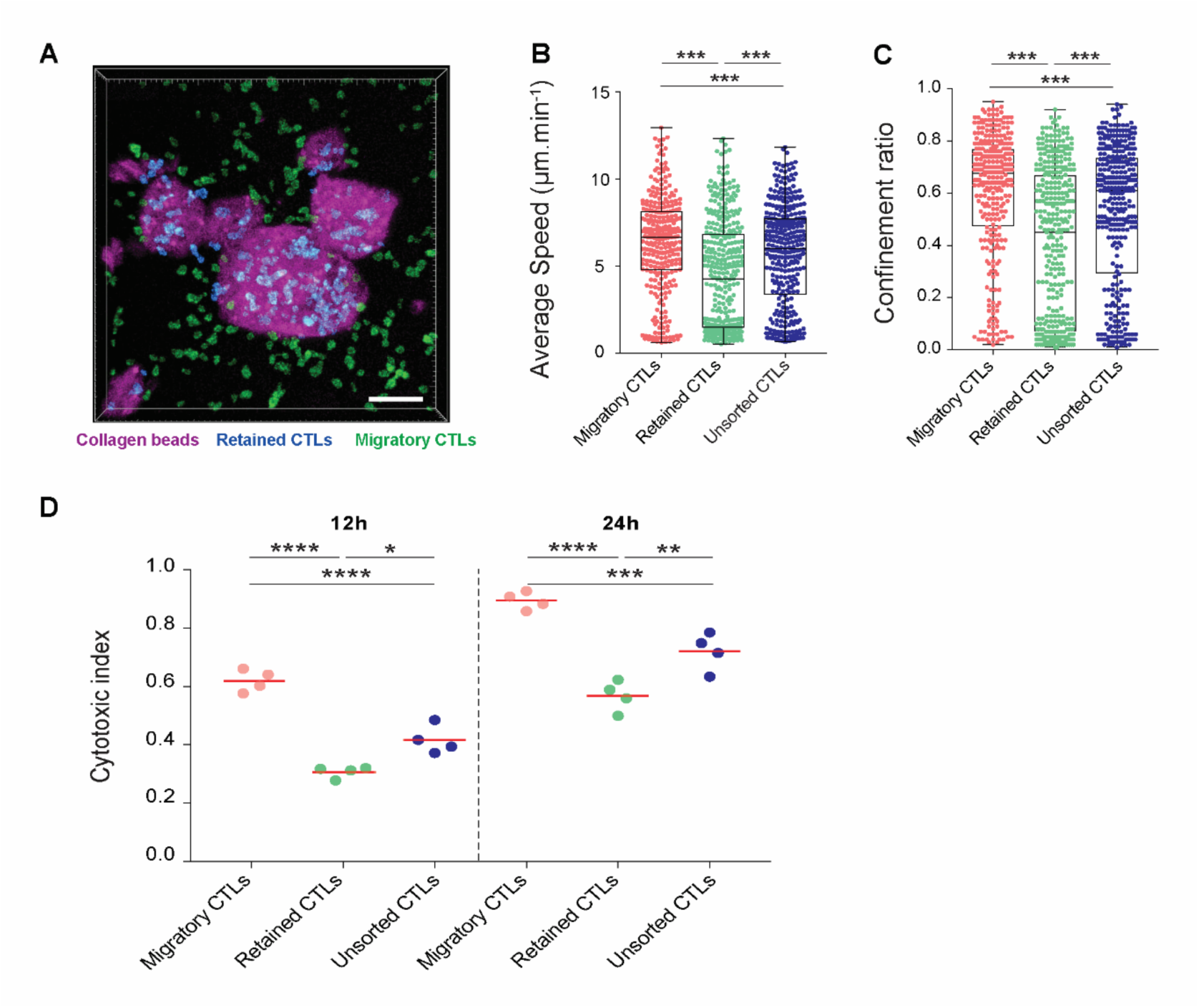
Migratory sorting and cytotoxic function of sorted T cells. (A) Confocal 3D image showing collagen beads (magenta) containing CTLs cultured for 2h in TCM under regular conditions. Using image analysis two T cell populations were identified. The CTLs that were retained in the collagen beads (blue cells) and the ones that migrated out (green cells). (B) Distribution of the average speed of migratory, retained or unsorted CTLs in 3D collagen matrices. Data points indicate individual tracks; box-whiskers: medians and quartiles from pooled data of 3 independent experiments; *** p<0.001 by Kruskal-Wallis test followed by Dunn’s multiple comparison test. (C) Distribution of the confinement ratio of migratory, retained or unsorted CTLs in 3D collagen matrices. Data points indicate individual tracks; box-whiskers: medians and quartiles from pooled data of 4 independent experiments; ***<0.001 by Kruskal-Wallis test followed by Dunn’s multiple comparison test. (D) Flow cytometry quantification of the cytotoxic index of migratory, retained or unsorted CTLs after 12 or 24h of incubation with target cells as is indicated. Red bars indicate the mean of 4 independent experiments. *<0.05, **<0.01, and ****<0.0001 by one way ANOVA followed by Tukey’s multiple comparisons test.

Finally, we tested the functional capability of the sorted T cells in terms of their ability to specifically kill target cancer cells, determined by a standard cytotoxicity assay based on flow cytometry analysis after 12 and 24 hours of co-incubation. Strikingly, it was observed that the faster migrating T cells had significantly higher killing efficiency as compared to both slow migratory and unsorted cells (Figure 6D). The cytotoxic index of the faster CTLs was about twofold higher as compared to slow migrating cells. This indicated the potential utility of this method for functional sorting of cytotoxic T cells in the context of immunotherapeutic applications. The potential reasons for such a correlation is discussed below.

Overall, our data thus demonstrates that we have established a 3D migration assay that can enrich fast migrating cells, which can then be recovered and tested for other correlative or causal factors.

## DISCUSSION

Single cell imaging and tracking has long established the heterogeneity of migration potential within a cell population. Besides optical characterisation, few tools exist that can phenotypically sort cells based on migration speed. Migration chromatography approaches are often limited by the complexity or the low yield of the devices that preclude subsequent testing of the sorted populations. For example, Chen et al., designed a microfluidic migration platform which can be used to profile chemotactic heterogeneity in cells at a single cell resolution. The migration in their work is however strictly driven by chemokine gradient and the process can yield only a limited number of cells (tens of cells) per experiment ^9^. In another method, phagocytosed fluorescent particles coated on the substrate are used as a measure of cell migration, this can be later concatenated with FACS to separate cells with different migration rates ^7^. While this method provides excellent single cell resolution in terms of describing migration ability of cells, the downstream analysis results may be significantly clouded by the presence of internalized particles. It is known that internalization of microplastics significantly alters cell physiology, and is known to elicit oxidative and inflammatory response in a dose dependent manner ^22^. Conversely, our approach adds cell sorting capabilities to the classical wound healing tests by creating layered substrates that can be physically detached from each other after migrations. The separation process is microscopic (tweezers or Sieve), in 2D and in 3D. The assay does not modify the classical conditions of migration typically used in wound healing assays. In particular, collective effects based on conditioning by exogeneous secretion or crowding effects can be readily controlled. The method here utilised collagen and hyaluronic acid which are both abundant in most human tissues. Further, like the 2D system this system is extendable to other matrix molecules which can be added to bulk HA gels to select for cells that migrate better in their presence. Furthermore, the system can be adapted for chemotactic migration by incorporation of immobilized chemokines and growth factors in the outer HA hydrogel.

The 2D or 3D implementation of the assay does not involve any obstacle as in other chromatography approaches. It relies on the ability of the fastest cells to emigrate from the initial colonies. Hence there is no intrinsic separation power of our technique since it depends on the collective behavior of the cells in the colony. The parameters of spacer thickness and cell density can be varied for different cell types. We found that for most cell lines an initial confinement on 300-500 μm diameter pits (respectively spheres) provide an adequate surface to perimeter ratio (respectively volume to surface ratio) to allow migratory cells to explore the colony boundaries and offer them the possibility to leave it. The sorting stringency can also be finetuned by modulating the time of migration, as was seen in case of HN cells, such that shorter incubation times stringently select more migratory cells. Hence the sorting itself is a consequence of the device and of the initial population. We here argue that the assays are thus physiologically relevant since cell migration or extravasation *in vivo* results from the intrinsic migratory potential of cells combined with their environmental conditions including neighbouring cells. Our functional assay explores the metastatic potential of a subpopulation of cells within a given mixed population.

Importantly, the 3D migration method opens avenues for migratory sorting of cells that do not migrate well on 2D substrates such as T cells. Here we demonstrate the possibility of effectively sorting fast migrating cytotoxic T cells using our method. Migratory sorting of T cells is of very high relevance because of their need to traverse large distances in tissues before they attack their targets ^23^. Killing of cancer cells in case of solid tumors involves the need to penetrate dense tissue before these cells can reach the core of the tumor. Strikingly, we report a strong correlation between migratory function and killing function of CTLs. It is beyond the scope of this paper to establish the molecular reasons why these two distinct properties of the T cells (cell migration and killing function) are actually coupled. We can however speculate that the correlation can arise from the orchestration of these two distinct mechanisms by actin regulation ^24,25^. The front-rear polarity development and persitence is largely governed by actin cytoskeleton dynamics ^26^. The killing potential of T cells is equivalently linked to the actin-based formation of immunological synapses ^27^. Our method thus may be of great importance to sort cells that would be able to migrate faster and better eliminate their target cancer cells in immunotherapy.

Finally, the methods we propose here are easily scalable due to the simplicity of fabrication. The multilayer pits are fabricated on large PDMS sheets, drilled by standard laser cutter and subsequently diced to the 6 well plate size. The 3D version relies on a standard emulsification method. We believe that the combination of the power of the sorting method and the ease of implementation is key for a direct adoption by other researchers and in clinical settings.

## MATERIALS AND METHODS

### Fabrication of layered PDMS micro-wells

Four PDMS sheets of thickness 125 or 250 µm were layered over each other, and through holes of diameter 500 µm were created in it using a laser cutting machine as depicted in Figure 1. The ash and debris from laser cutting were cleaned by sonication in isopropanol/water mixture (1:1) for 10 mins. The device was rinsed several times with water, followed by drying using pressurized nitrogen. Finally, the PDMS layered device with through holes was layered over another PDMS sheet (125 µm). The large array was then cut into sizes as per experimental requirements. The micro-well arrays were attached to tissue culture plates using Norland optical adhesive (NOA) followed by UV curing.

### Cell sorting using PDMS microwells

The PDMS micro-wells were UV sterilized and coated with fibronectin (50 µg/mL) (Roche Lifescience). For cell seeding, 500 µl of cell suspension (~ 6 −7 x10^5^ cells/ml) was added gently onto 6×6 wells/platform. Once the cells settle down inside the wells by gravity, the excess cells on the protecting layer are removed by giving a gentle wash to the PDMS device. The protecting layer is then peeled off with the help of sharp forceps to remove any cells on the top of the device. Finally, enough media is added to completely submerge the device. The cells then migrate during the incubation period of 3-5 days. The collecting layer with the migrated cells can then be peeled off, as shown in Figure 1. The cells can thereafter be isolated by trypsinization from PDMS sheets and can then be cultured further or used to any other analysis.

### EMT scoring

Based on a previous study ^28^, EMT scoring was performed by analyzing gene expression data base on micro-array analysis of a subset of cancer related genes to score the different population in terms of their EMT status. To compute the EMT score in cell lines, a similar approach that used in ssGSEA ^28^ was adopted. Empirical cumulative distribution function (ECDF) was estimated for Epithelial and Mesenchymal gene sets. The 2KS test was employed to compute the difference between the Mesenchymal ECDF (ECDF_Mes_)and the epithelial ECDF (ECDF_Epi_). The 2KS score was then taken as the EMT score. A cell line with a positive EMT score exhibits a more “mesenchymal-like” phenotype, whereas a negative EMT score reflected a more “epithelial-like” phenotype.

### Ephrin clustering assay

Ephrin clustering assay was performed as reported previously ^18^. Briefly, 9 chambers (3 x 3 mm) were formed by placing a silicon gasket on a clean, hydrophilic glass coverslip. Supported lipid bilayers with 96 mol% DOPC (Avanti Polar Lipids) and 4% DOGS-NTA-Ni (Avanti Polar Lipids) were then deposited using Small Unilamellar Vesicle (SUV) deposition. Polyhis-ephrinA1 labeled with Alexa 568 was then allowed to bind NTA lipid moieties at saturating densities. The labelled bilayers were then washed. Cells were seeded on the wells and allowed to attach on the bilayer surface. The samples were fixed after one hour and imaged using a fluorescence microscope.

### Sorting of cancer cells using hierarchical hydrogels

For fabrication of collagen micro-gels, cells (either MDA MB 231 GFP or a mixture of MDA MB 231 GFP and MCF 7 at a ratio of 1:1) were suspended in cold neutralized bovine collagen solution (2.5 mg/ml) (Thermo Fisher Scientific). This cell suspension was then added to pre-warmed (37 ̊C) mineral oil with 0.2% Span 80 while slowly stirring the oil using a magnetic stirrer. The emulsion was allowed to stir for 45 minutes. Thereafter, the emulsion was centrifuged at 1500 rpm for 5 minutes to separate micro-hydrogels from the oil phase. The pellet was washed thrice with complete cell culture medium (DMEM with 10% FBS) to remove traces of oil and surfactant. Finally, the gels were suspended in medium and cultured under regular cell culture conditions.

The fabricated micro-hydrogels after culture for 24 hours were suspended in hydrogel precursor solution containing Hyaluronic acid-tyramine (5 mg/ml) (LifeCore Biomedical), collagen (0.5 mg/ml) and horseradish peroxidase (2.5 U/ml) solution in phosphate buffered saline. Droplets of 100 µl of this suspension were then deposited in a 12 well plate. Thereafter, hydrogen peroxide was added to the droplets at a final concentration of 3 mM and mixed rapidly to enable cross-linking and gelation (< 1 minute). The gels were then incubated in complete cell culture medium at 37 ̊C and 5% CO2.

After 7 days of in vitro culture the samples were digested enzymatically using hyaluronidase (1 mg/ml in serum free DMEM media) (Sigma Aldrich) for 3 hours at 37 ̊C. This solution was then passed through a 40 µm cell strainer to separate single cells migrated out into bulk hydrogels from the cells retained in the micro-gels. The filtrate and the retentate containing the migrated and non-migrated cells, respectively, were plated on regular cell culture dishes to enable culture and analysis of the two sub-populations.

### T Cell sorting using collagen microbeads

T cells were isolated from OT-1 x Lifeact-EGFP mice spleens. To generate effector CD8^+^ T cells, splenocytes were stimulated with SIINFEKL peptide for 4h in T cell medium (TCM) and were then cultured with IL-2 for 6-7 days. For generation of collagen micro-gels 1 × 10^6^ effector CTLs were suspended in neutralized ice-cold rat-tail collagen I solution (50 µl at 1.75-2 mg/ml) (Corning). The addition of Alexa Fluor 647 succinimidyl ester (1µl) allowed the visualization of the collagen particles during imaging. This cell suspension was then added to pre-warmed (37 ̊C) mineral oil with 0.2% Span 80 while slowly stirring the oil using a magnetic stirrer. The solution was stirred for 30 min allowing the formation of collagen micro-gels. Thereafter, the emulsion was centrifuged at 1500 rpm for 5 minutes to separate micro-hydrogels from the oil phase. The pellet was washed 3 times with TCM to discard traces of oil and surfactant. The collagen beads were incubated in 24 well plates for 2 h under regular culture conditions to allow T cells to migrate out of the beads. The cell suspension that contained the micro-gels was passed through a 70 µm cell strainer separating the CTLs in suspension (migratory CTLs) from those retained in the micro-gels. Then the collagen micro-gels were treated with collagenase IV (0.5 mg/ml for 30 minutes) (Sigma Aldrich) to isolate the non-migrated cells (retained CTLs). To account for the effects of collagenase, unsorted CTLs (mix population with both migratory and retained cells) and the migratory CTL population were also treated with collagenase IV.

To validate our functional sorting, the isolated CTL populations were re-embedded in collagen matrices and live imaged for 1 h to quantify migration parameters. Individual cells were then tracked in 3D and their average speeds and confinement ratios were measured.^21^

### Cytotoxic activity of T cells

Cytotoxicity assays were performed using flow cytometry based on the ratio between live target and non-target cells. A total number of 1×10^5^ OT-1 T cell was mixed with cognate (+SIINFEKL) and non-cognate (-SIINFEKL) EL4 tumour cells (ATCC: TIB-39) at a ratio of 1:4:4. The cognate and non–cognate cells were differentially labelled with cell tracker® dyes and thus could be counted using flow cytometry analysis. Following exclusion of death cells (DAPI positive cells), the extent of reduction in the ratio of target to non-target cells with respect to a control mix lacking T cells was used as the measure of killing capacity of T cells.

